# Cryo-EM structures reveal a multistep mechanism of Hsp90 activation by co-chaperone Aha1

**DOI:** 10.1101/2020.06.30.180695

**Authors:** Yanxin Liu, Ming Sun, Alexander G. Myasnikov, Daniel Elnatan, Nicolas Delaeter, Michael Nguyenquang, David A. Agard

## Abstract

Hsp90 is a ubiquitous molecular chaperone that facilitates the folding and maturation of hundreds of cellular “client” proteins. The ATP-driven client maturation process is regulated by a large number of co-chaperones. Among them, Aha1 is the most potent activator of Hsp90 ATPase activity and thus dramatically affects Hsp90’s client proteins. To understand the Aha1 activation mechanism, we determined full-length Hsp90:Aha1 structures in six different states by cryo-electron microscopy, including nucleotide-free semi-closed, nucleotide-bound pre-hydrolysis, and semi-hydrolyzed states. Our structures demonstrate that the two Aha1 domains can each interact with Hsp90 in two different modes, uncovering a complex multistep activation mechanism. The results show that Aha1 accelerates the chemical step of ATP hydrolysis like a conventional enzyme, but most unusually, catalyzes the rate-limiting large-scale conformational changes of Hsp90 fundamentally required for ATP hydrolysis. Our work provides a structural framework to guide small molecule development targeting this critical modulator of client protein maturation.

## Introduction

Heat Shock Protein 90 (Hsp90) is one of the most abundant proteins in the cell. It is a molecular chaperone that hydrolyzes ATP to assist the folding and maturation of hundreds of “client” proteins^1,2^. These client proteins are highly enriched for signaling and regulatory proteins such as kinases, transcription factors, and E3 ligases^3^. Hsp90 has been increasingly recognized as a regulatory hub for maintaining protein homeostasis and cellular function^1,4^. As an important component of the stress response machinery, Hsp90 plays an essential role in many diseases including cancer^5,6^, infectious diseases^7,8^, and neurodegenerative diseases^9^ and has been exploited as a potential therapeutic target^10–12^.

Hsp90’s diverse cellular functions are supported and regulated by a group of more than 20 co-chaperones^13,14^. Through sequential interaction with Hsp90, co-chaperones modulate the Hsp90 conformational cycle and facilitate client loading, remodeling, and release. A key member in the co-chaperone family is the Activator of Hsp90 ATPase homolog 1 (Aha1), which is the most potent Hsp90 activator known to date^15^. The 39-kDa Aha1 has two domains, an N-terminal domain (Aha1_NTD_) and a C-terminal domain (Aha1_CTD_), connected by a 60-residue flexible linker. However, the molecular mechanism by which Aha1 regulates Hsp90 remains elusive.

Hsp90 functions as a homodimer and has a modular structure. Each protomer consists of three domains: an ATP-binding N-terminal domain (Hsp90_NTD_), a middle domain (Hsp90_MD_) that participates in co-chaperone and client binding, and a C-terminal domain (Hsp90_CTD_) responsible for client binding and dimerization. These three domains can move relative to each other providing Hsp90 with a high degree of conformational flexibility^16^. Although required for client maturation, Hsp90’s intrinsic ATPase activity is rather slow (1 ATP/min for yeast Hsp90 and 0.1 ATP/min for human Hsp90)^17–19^. The rate limiting step in the reaction is Hsp90 dimer closure, which is a prerequisite for ATP hydrolysis. As shown schematically in Fig.1A, the dimer closure represents a large energy barrier in the reaction pathway and requires a series of events to take place, including ATP binding, lid closure, Hsp90_NTD_ rotation, and Hsp90_NTD_ dimerization^16^. The co-chaperone Aha1 dramatically enhances Hsp90 ATPase activity by accelerating the Hsp90 dimer closure rate^19,20^. Once the Hsp90 dimer closes, ATP is readily hydrolyzed.

**Fig. 1.**
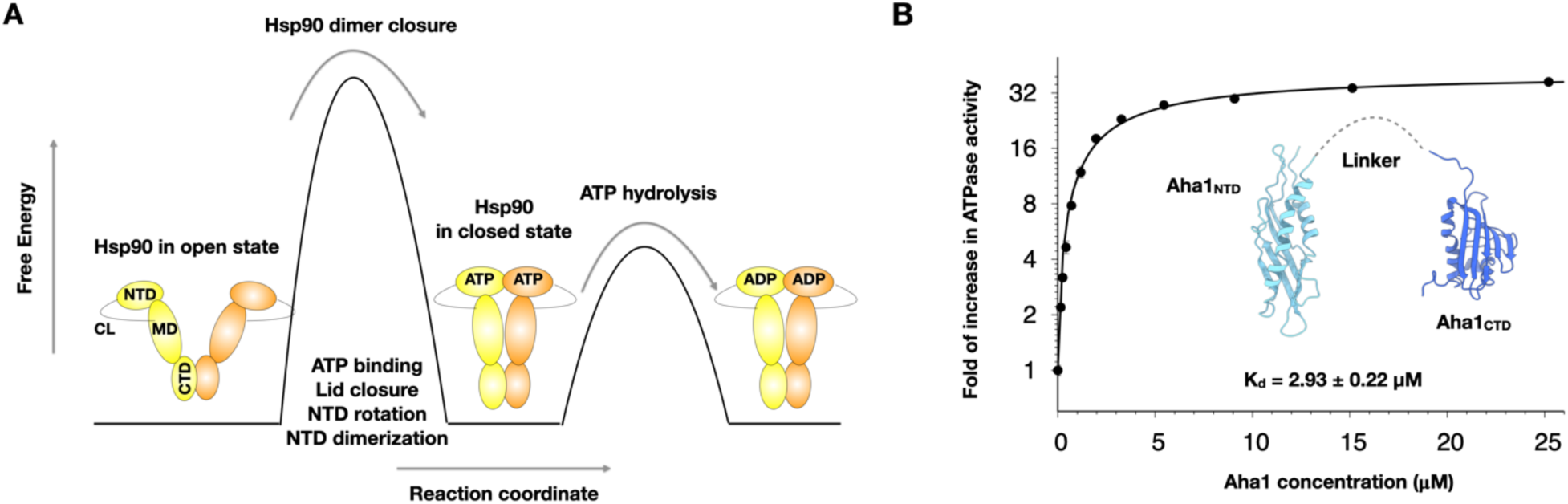
Effect of Aha1 on Hsp90 ATPase activity. (A) Hsp90 conformational steps leading to ATP hydrolysis. The Hsp90 dimer closure is the rate-limiting step. (B) Aha1 substantially accelerates Hsp90 (yeast Hsc82) ATPase activity. The full-length Aha1 structure is not known. The individual domain structures of Aha1 are shown. Aha1_NTD_ is colored cyan and Aha1_CTD_ is colored blue.

A fragment-based crystal structure of Hsp90_MD_ in complex with Aha1_NTD_ was reported previously providing a model for Aha1 recruitment to Hsp90 but little information on how it could modulate the Hsp90 conformational cycle^21^. The interactions between Hsp90 and Aha1 were also probed by NMR^22–24^, which were restricted to individual domains or specific isoleucine labeling. By working with full-length Hsp90 and Aha1, we determined 7 atomic structures for Hsp90 complexed with Aha1 representing 6 different states using single particle cryo-electron microscopy (cryo-EM). Many of the binding modes observed span Hsp90 domains or protomers and thus could never have been observed using domain fragments. These structures with Hsp90 in either nucleotide-free (apo) or nucleotide-bound states (AMPPNP, ATPγS, or ADP) reveal that each Aha1 domain can bind to Hsp90 in two different ways. As a result, Aha1 can stabilize a semi-closed Hsp90 apo state, as well as fully closed pre-hydrolysis and semi-hydrolyzed nucleotide bound states. Together they suggest a multistep mechanism through which Aha1 accelerates Hsp90 activity both conformationally and catalytically. These results provide an atomistic framework to guide small molecule development targeting the interactions between Aha1 and Hsp90 as a means to modulate the maturation of Hsp90-dependent client proteins.

### Aha1 accelerates yeast Hsp90 ATPase activity

Both yeast Hsp90 and Aha1 were recombinantly expressed and purified from *E. coli*. We chose to work on the dominant, constitutively expressed Hsp90 isoform (Hsc82) from *S. cerevisiae* (referred to as Hsp90 for simplicity unless noted). Similar to the stress-induced Hsp82 isoform^21^, we observed a 35-fold increase in Hsc82 ATPase activity under saturating Aha1 concentrations (Fig.1B). The calculated EC_50_ was ∼2.9 μM, indicating an effective affinity comparable to that reported for Hsp82 or measured for other co-chaperones^20^.

Although the structure of full-length Aha1 has not been determined, the structures of its two individual domains are known. The atomic structure of yeast Aha1_NTD_ in complex with yeast Hsp82_NTD_ was solved by X-ray crystallography (PDB: 1USV)^21^ and the human Aha1_CTD_ structure was determined by NMR (PDB: 1×53). A flexible 60-residue linker connecting Aha1_NTD_ and Aha1_CTD_ was not resolved. The intrinsic flexibility of Aha1 and Hsp90, as well as varied binding stoichiometry^20^, have hindered the structural understanding of the complex. We decided to solve the structures for Hsp90:Aha1 complex using cryo-EM, which has the ability to sort out both conformational and compositional heterogeneity through computational image analysis.

### Aha1 bridges apo Hsp90 to induce a semi-closed state

Hsp90 adopts an open V-shape in apo state and will slowly convert to a closed state in the presence of non-hydrolyzable analogs of ATP^25^. Previous results from analytical ultracentrifugation showed that Aha1 can bind to Hsp90 in both the open and closed states^20^. We began by exploring what Hsp90 state is induced and stabilized by Aha1 in the absence of nucleotides. The binding affinity between Aha1 and Hsp90 is weaker in the apo state (∼1.2 μM) than in the closed state (∼0.16 μM) as determined previously by surface plasmon resonance^20^. To overcome the challenge of weak and transient association, we mixed 10 μM Aha1 and 2 μM Hsp90 to achieve an oversaturated condition and carried out a single particle cryo-EM investigation. The 39-kDa Aha1 is small enough so that the excess does not significantly contribute to micrograph background.

We determined the cryo-EM structure of Hsp90/Aha1 complex in the apo state to 3.8 Å resolution (Fig.2A, Fig. S1). The structure had C2 symmetry and consisted of two full-length Aha1 molecules and one Hsp90 dimer (Hsp90^apo^-Aha1_NCNC_). The observed stoichiometry of 2:1 (Aha1:Hsp90 dimer) likely resulted from the high Aha1 concentrations used for cryo-EM. Under physiological conditions with much less Aha1 than Hsp90, one Aha1 binding per Hsp90 dimer is the most likely scenario. The binding between Aha1_NTD_ and Hsp90_MD_ closely resembles the previously determined fragment crystal structure (Fig, 2B), which was suggested to represent a recruitment step in Aha1 binding to Hsp90^21^. As shown in Fig. 2A, while Aha1_NTD_ binds to Hsp90_MD_ from one Hsp90 protomer with a buried surface area (BSA) of 755 Å^2^, the Aha1_CTD_ binds to the Hsp90_MD_ of both protomers with BSA of 293 Å^2^ and 586 Å^2^ respectively and the Hsp90_CTD_ amphipathic helix from one of the protomers with BSA of 562 Å^2^. The Hsp90_CTD_ amphipathic helix had previously been observed to bind client proteins in the closed state^26^, but here is repurposed for co-chaperone binding. The cross-protomer interaction facilitated by Aha1_CTD_ coupled with the direct interaction between Aha1_NTD_ and Aha1_CTD_ (BSA=421 Å^2^) lead to a partial closing of the two Hsp90 protomers, contrasting with the widely opened resting Hsp90^apo^ state. Deleting either Aha1 domain almost completely abolished its ability to accelerate Hsp90 ATPase, although supplying them *in trans* restores partial activity (Fig. S2). Thus, through multiple interaction surfaces the full-length Aha1 serves as a bridge that connects the two Hsp90 protomers and induces a semi-closed Hsp90 that is on-pathway to full closure. Compared to the open state of apo HtpG crystal structure^25^, the Hsp90_MD_ rotates 8° relative to the Hsp90_CTD_ (Fig. 2C left, Movie S1). However, even in the presence of Aha1 Hsp90^apo^ does not fully close. As shown in Fig. 2C (right), the Hsp90_MD_ would need to further rotate by 5° to match with closed state of yeast Hsp82 (Fig 2A) as stabilized by nucleotide AMPPNP and co-chaperone p23 in the crystal structure^27^ (Movie S1).

**Fig. 2.**
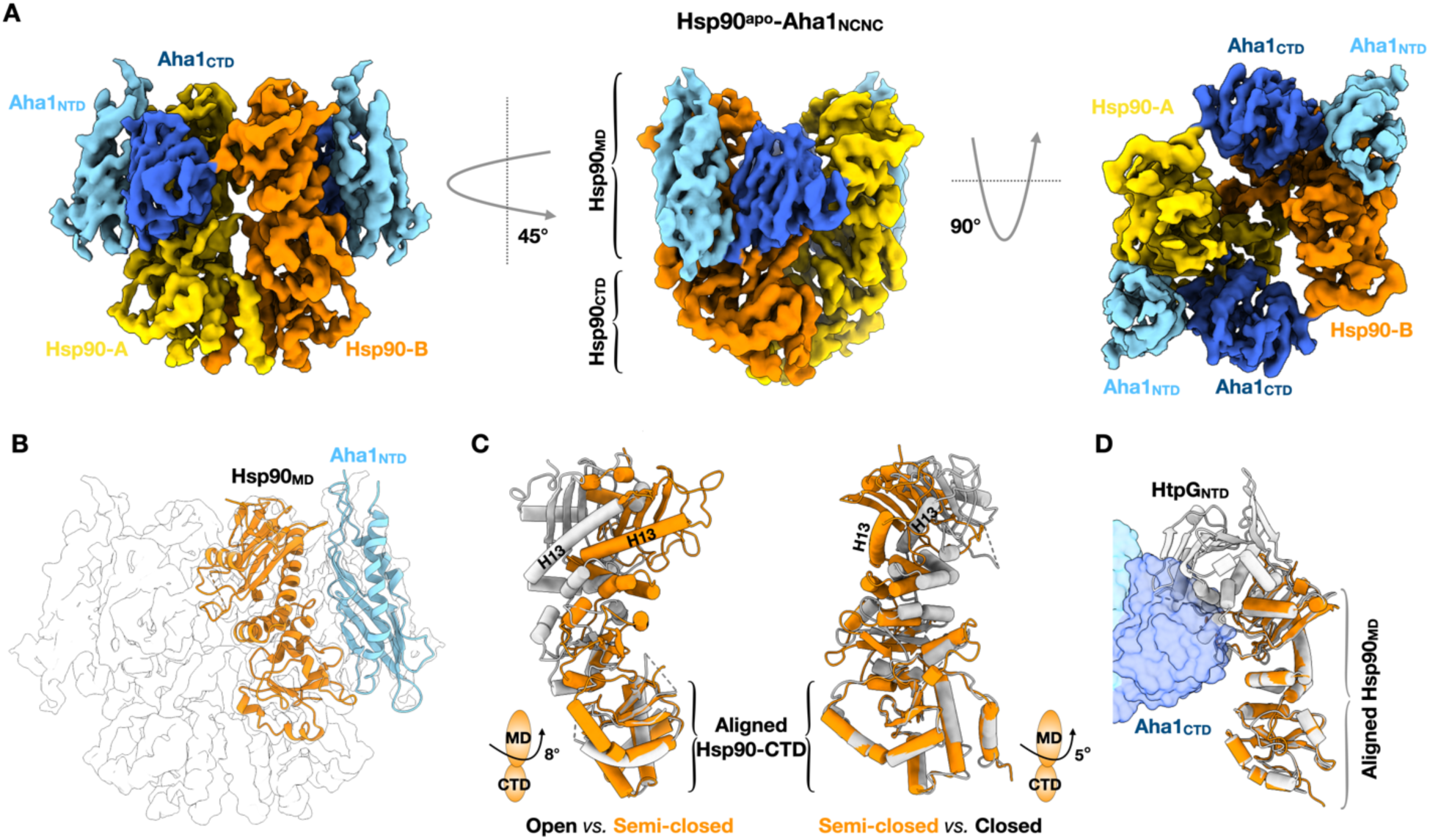
Aha1 induces a semi-closed state of Hsp90 in the absence of nucleotides. (A) Cryo-EM structure of Hsp90 dimer in complex with two Aha1 in the absence of nucleotide. (B) The interacting mode between Aha1_NTD_ and Hsp90_MD_ is consistent with previously determined fragment-based crystal structure (PDB: 1USV). (C) The Aha1-stabilized state shown in orange represents a semi-closed intermediate state between the fully open state of bacterial Hsp90 (HtpG, PDB: 2IOQ, gray on the left) and the fully closed state of yeast Hsp90 (Hsp82, PDB: 2CG9, gray on the right). (D) The Hsp90_NTD_ in the fully open state of HtpG (PDB: 2IOQ) sterically clashes with Aha1_CTD_ (blue surface). This clash results in the undocking and the missing density for the Hsp90_NTD_ in our cryo-EM structure shown in (A).

### Aha1 binding undocks Hsp90 NTD in the apo state

A striking feature in the reconstruction of the Hsp90^apo^:Aha1 complex is that all density for the Hsp90_NTD_ was missing (even when low-pass filtered), despite the sample being full-length. We hypothesize that the Hsp90_NTD_ as a rigid domain, is still connected with Hsp90_MD_ through the long charged linker (CL) but becomes disordered relative to the rest of the Hsp90. As shown in Fig. 2D, when the apo structure of the bacterial Hsp90 (HtpG) is superimposed on our Aha1-bound semi-closed state, the HtpG_NTD_ has significant steric clash with the Aha1_CTD_. From this, we conclude that binding of Aha1 onto Hsp90^apo^ is sterically incompatible with the resting conformation of Hsp90_NTD_ and undocks the Hsp90_NTD_. Hsp90_NTD_ was observed to easily undock using optical tweezers^28^. Subsequent redocking in a rotated conformation would then be on-pathway to closure and Hsp90_NTD_ dimerization^29,30^.

### Aha1 stabilizes AMPPNP-bound closed states of Hsp90

Although Aha1 binding induces semi-closure of apo Hsp90, nucleotide binding is required for Hsp90 to fully close in order to reach the ATP hydrolysis competent state. To investigate the effect of Aha1 on the Hsp90 closed state, we determined the cryo-EM structure of Hsp90:Aha1 complex in the presence of the non-hydrolyzable ATP analog AMPPNP (Fig. 3A-D). The cryo-EM datasets were collected at Aha1:Hsp90 dimer concentrations ratios of 0:4, 4:4, 5:2, and 40:10 (all in μM).

**Fig. 3.**
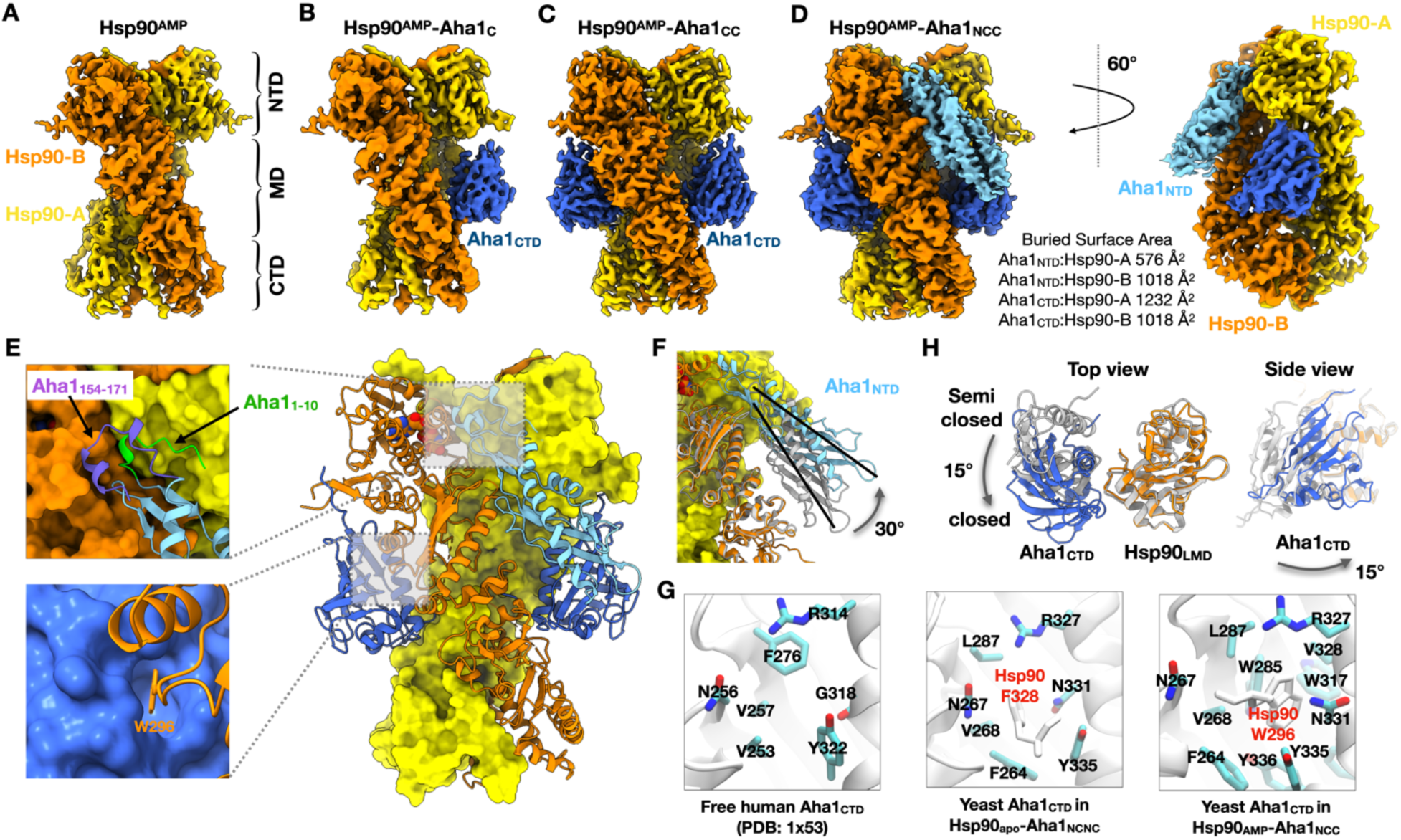
Cryo-EM structures of Hsp90 dimer in the presence of AMPPNP (A) without Aha1. (B) with Aha1_CTD_. (C) with two Aha1_CTD_. (D) with Aha1_NTD_ and two Aha1_CTD_. (E) Atomic model of Hsp90 dimer with Aha1_NTD_ and two Aha1_CTD_ derived from EM density shown in (D). Upper left inset shows the interface between Hsp90_NTD_ and Aha1_NTD_. The first 10 residues from Aha1_NTD_ are colored green and last 18 residues (I154-K171) from Aha1_NTD_ are colored purple. These 28 residues are unstructured in the previously determined fragment-based crystal structure (PDB:1USV). Lower left inset shows the interface between Hsp90_MD_ and Aha1_CTD_. The key event is W296 from Hsp90_MD_ inserts into Aha1_CTD_ through an induced fit mechanism. (F) Aha1_NTD_ (cyan) alters its binding interface with Hsp90 by tilting up 30° when interacting with the closed state Hsp90 compared to the semi-closed state of Hsp90 shown in Fig. 2 (gray). (G) Aha1_CTD_ switches its binding interface with Hsp90 in different states. (H) Aha1_CTD_ (blue) utilizes a different interacting interface with Hsp90_MD_ when interacting with the closed state Hsp90 compared to the semi-closed state of Hsp90 shown in Fig. 2 (gray). Only Aha1_CTD_ and Hsp90 large middle domain (LMD) are shown in top view for clarity.

In the absence of Aha1, the structure of AMPPNP-bound Hsp90 by itself was determined at 3.7 Å resolution (Hsp90^AMP^, Fig. 3A, Fig. S3A). Two structures of full-length cytosolic Hsp90 were solved previously. The first one is the yeast Hsp82 in complex with co-chaperone p23 by X-ray crystallography^27^. The second one is the human Hsp90β in complex with the Cdc37 co-chaperone and client kinase Cdk4^31^. Our structure represents the first for a standalone full-length cytosolic Hsp90. It is almost identical to the crystal structure of yeast Hsp82 stabilized by p23 (RMSD: 0.925 Å). In contrast to the Hsp82 crystal structure in which the charged linker was truncated, our cryo-EM map clearly shows density for the charged linker when the map is low-pass filtered to 5 Å (Fig. S3B). However, the flexible nature of the charged linker limited the ability to model most of the linker.

At a 4:4 (Aha1:Hsp90) µM ratio in the presence of AMPPNP three Aha1 binding modes exist in equilibrium: Hsp90 alone (low quality and not shown), Hsp90 binding with one Aha1_CTD_ reconstructed at 3.7 Å resolution (Hsp90^AMP^-Aha1_C_, Fig. 3B, Fig. S4A), Hsp90 binding with two Aha1_CTD_s. Although full-length Aha1 was used throughout, the Aha1_NTD_ was not visible, presumably due to flexibility. Increasing Aha1 concentration (40:10 both in µM) shifted the equilibrium and revealed two Hsp90 closed state reconstructions, one with two Aha1_CTD_s and the other with one Aha1_NTD_ and two Aha1_CTD_s. As the Hsp90^AMP^-Aha1_CC_ reconstructions from all datasets were the same, those particles were combined to yield a final reconstruction at 2.74 Å resolution (Hsp90^AMP^-Aha1_CC_, Fig. 3C, Fig. S4B). The Hsp90^AMP^-Aha1_NCC_ reconstruction was refined to 3.15 Å resolution (Hsp90^AMP^-Aha1_NCC_, Fig. 3D, Fig. S4C). Owing to the improved map quality over the crystal structure of p23 stabilized Hsp82^27^, we refined the atomic model of Hsc82 by fixing registry errors in the lid helix region and the β-hairpin connected to the charged linker. The Hsc82 conformation is the same in all four of these AMPPNP cryo-EM structures (Fig. 3A-D) and is also very similar to the Hsp82 crystal structure with RMSD being 1.056 Å, 1.061 Å, 1.069 Å, 1.083 Å respectively. Therefore, we focused our structural analysis on the different binding modes between Aha1 and Hsp90.

The atomic model derived from the density map of Hsp90^AMP^-Aha1_NCC_ is shown in Fig. 3E. Compared to the apo state complex, the Aha1_NTD_ tilts up by 30° (Fig 3F, Movie S2) breaking the interface with the Hsp90_MD_ (Fig 2A,B) and then forming a new interface that spans across the Hsp90_NTD_ dimer (Fig. 3E upper left inset). The BSA between Aha1_NTD_ and Hsp90 increased from 755 Å^2^ to 1562 Å^2^, suggesting much tighter binding to this state. Aha1_NTD_ residues 1-10 (green in Fig. 3E upper left inset) and residues 154-171 (purple Fig. 3E upper left inset) become ordered upon forming this new Hsp90_NTD_ interface. In contrast, these residues were unstructured in Hsp90^apo^-Aha1_NCNC_ state and the Hsp90_MD_-Aha1_NTD_ crystal structure (Fig. 2). Supporting the functional relevance of the Aha1_1-10_ interaction, deletion of the N-terminal 11 amino acids harboring the conserved NxNNWHW motif in Aha1 reduced its ATPase stimulation effect by 2.8 fold and abolished its ability to rescue a temperature sensitive yeast strain with harboring the Hsc82_S25P_ mutation^32^. The Aha1_154-171_ forms a short helix hairpin that is in close contact with the Hsp90 lid region and may accelerate ATP hydrolysis rate as discussed below in the context of ATPγS. The observed Aha1_NTD_ movement likely takes place after the Hsp90_NTD_s dimerize as only then would the full interaction surface be available.

As Hsp90 transitions from the apo state to the fully closed state, the interaction mode between Aha1_CTD_ and Hsp90_MD_ also changes. In both states, the surface patch on Aha1_CTD_ formed by residues F264, N267, V268, L287, R327, N331, and Y335 interacts with the Hsp90_MD_ (Fig. 3G). However, the region on Hsp90_MD_ switches from the amphipathic loop (residues 320 to 336, also known as the Src-loop) to the loop (residues 293 to 300) connecting helix 11 and strand 10. Both loops were previously reported to interact with client proteins^33,34^, again revealing how client binding residues are repurposed for co-chaperone binding. This switch is achieved by a 15° counterclockwise rotation (viewed from the top) of Aha1_CTD_ relative to the Hsp90_MD_ (Fig 3H, Movie S3). As shown in Fig. 3E (lower inset), when bound to a closed Hsp90 a deep hydrophobic pocket opens in Aha1 to accommodate W296 from the Hsp90_MD_. A similar but much shallower pocket was observed when Aha1 binds apo Hsp90 (the semi-closed state in Fig. 2A) and was occupied by F328 on the Src-loop from the Hsp90_MD._ In the NMR structure of the free form of Aha1_CTD_, the patch forming residues mentioned above move close to each other to further diminish the pocket (Fig. S5). Mutating W296 on Hsp90 to either Ala or Gly reduces the Aha1 ATPase activation by 6-fold (Fig. S6) indicating the importance of this interaction. A recent small molecule screen identified compound SEW84 which completely inhibits Aha1 stimulation of the Hsp90 ATPase^35^. NMR chemical shift perturbations induced by SEW84 binding to the Aha1_CTD_ revealed the same residues that form the hydrophobic pocket occupied by F328 and W296 in our cryo-EM structures. We thus conclude that SEW84 competitively binds to the same pocket on Aha1_CTD_ as does Hsp90 F328 and W296, abolishing the Aha1 interaction and Hsp90 ATPase stimulation. Our structures also revealed that in a different binding mode Aha1_CTD_ bridges the two Hsp90 protomers, further stabilizing the closed state of the Hsp90 via interactions with Hsp90_NTD_ and Hsp90_MD_ from one protomer, as well as Hsp90_MD_ and Hsp90_CTD_ from the other protomer (Fig. 3D). However, unlike in the apo state, the Aha1_NTD_ and Aha1_CTD_ do not interact with each other in the Hsp90 closed state (Fig. 3D).

### Aha1 catalyzes sequential Hsp90 ATP hydrolysis

The cryo-EM structures described above revealed the effect of Aha1 on the Hsp90 open-closed conformational transition. We next investigated whether there was any effect of Aha1 on the chemical step of Hsp90 ATP hydrolysis. To do this, we repeated the structural analysis using the slowly hydrolyzable ATPγS instead of the non-hydrolysable AMPPNP. The hydrolysis rate of ATPγS is sufficiently slow to accumulate Hsp90 closed states during the timeframe of cryo-EM sample preparation. However, as the hydrolysis rate will be accelerated by Aha1, a sub population of the complexes captured by cryo-EM will be in a post-hydrolysis state. For this, 40 μM Aha1 was incubated with 10 μM Hsp90 for 30 mins in the presence of ATPγS. This ratio and concentration of Aha1 and Hsp90 were shown to yield the best structure in the presence of AMPPNP (Fig. 3C-D).

Structures for two different ATPγS bound Hsp90:Aha1 complexes were obtained. One has only two Aha1_CTD_s bound to the closed state of Hsp90 (Hsp90^ATPγS^-Aha1_CC_, Fig. 4A), whereas the other has additional density for the Aha1_NTD_ (Fig. 4B). The overall structure and the binding modes are identical to the ones solved using the same protein concentrations with AMPPNP. Using a smaller pixel size (0.822 Å) to achieve better resolution, the two structures were refined to 2.71 Å and 2.83 Å respectively (Fig. S7). In the Hsp90^ATPγS^-Aha1_CC_ structure where the Aha1_NTD_ was not visible, both ATP binding pockets were clearly occupied by ATPγS (Fig. 4A, inset). This structure thus corresponds to a pre-hydrolysis state of Hsp90. The surprise came with the second structure that also has the Aha1_NTD_ bound to dimerized Hsp90_NTD_. While the Hsp90 are symmetric, the Aha1 binding is asymmetric with a single Aha1_NTD_ on one side of the complex. The consequence of this broken symmetry is an asymmetric hydrolysis state in the two Hsp90 protomers. In the view shown, the Hsp90 protomer on the right side of Aha1_NTD_ still processes an ATPγS (right inset, Fig 4B), whereas ATPγS has clearly been hydrolyzed to ADP in the other protomer (left inset, Fig 4B). From this, we conclude that a semi-hydrolyzed Hsp90 state was captured (Hsp90^ATPγS/ADP^-Aha1_NCC_), also establishing the time sequence for these two structures. Nucleotide binding first induces the closure of the Hsp90 dimer with Aha1_CTD_ bound (Fig. 4A). Subsequent Aha1_NTD_ binding to Hsp90_NTD_ then catalyzes the ATP hydrolysis in one of the protomers resulting in a semi-hydrolyzed state (Fig. 4B). The accelerated hydrolysis on one protomer likely results from the direct interaction between the Aha1_NTD_ and the lid region covering the bound nucleotide in that protomer (Fig. S8). The interaction stabilizes the lid of the Hsp90 ATP binding pocket, providing a preferred local environment for ATP hydrolysis. In addition, once Aha1 has bound, which Hsp90 protomer hydrolyzes ATP first becomes deterministic. This could have important consequences for the progression of the Hsp90 cycle in the context of bound clients or other co-chaperones which themselves could either induce or respond to an asymmetry.

**Fig. 4.**
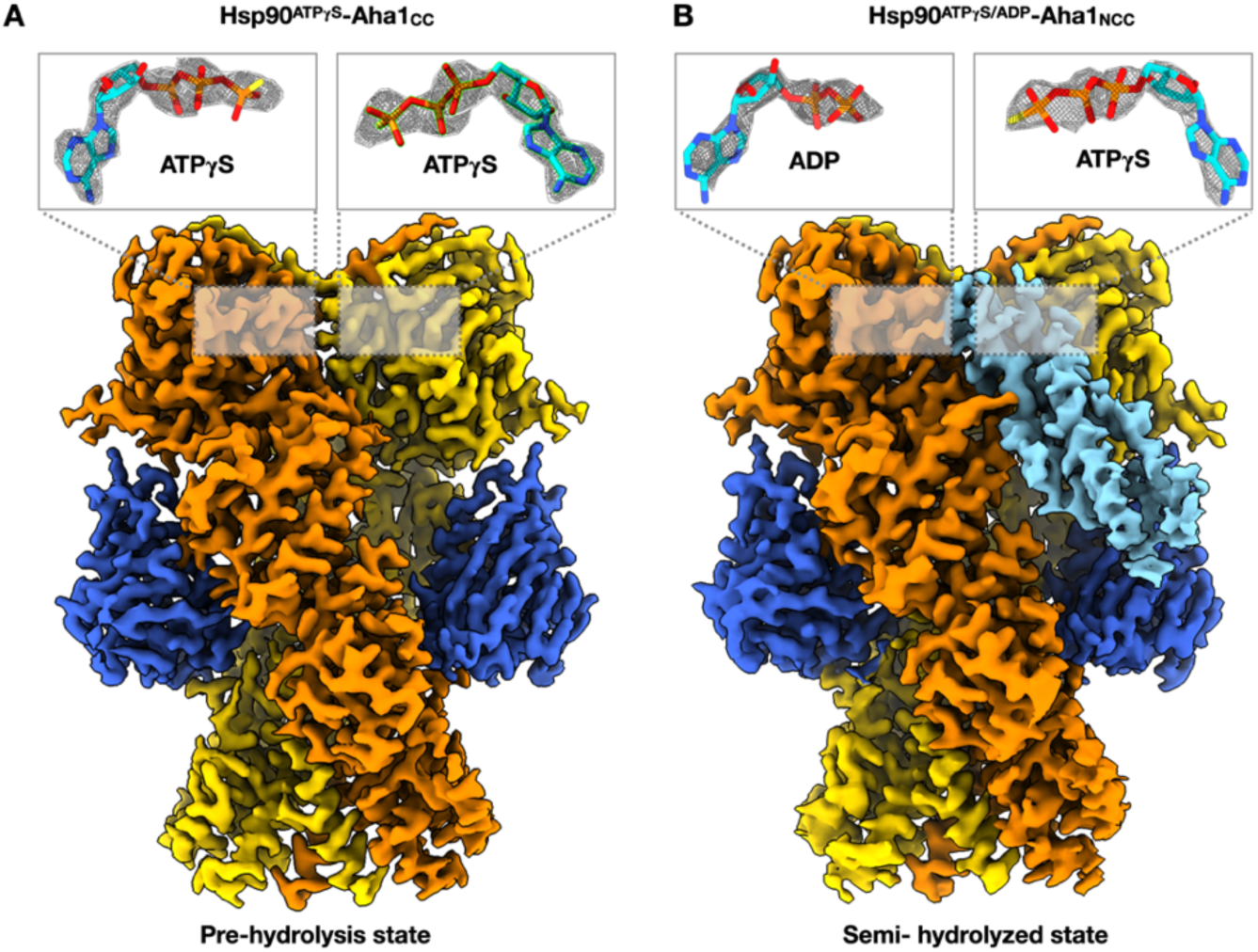
Cryo-EM structure of Hsp90 in complex with Aha1 in the presence of ATPγS. (A) Hsp90 dimer binding with two Aha1_CTD_. Non-hydrolyzed ATPγS densities are clearly visible in the nucleotide binding pocket of Hsp90. (B) Hsp90 dimer binding with one Aha1_NTD_ and two Aha1_CTD_. The Aha1_NTD_ binding results in ATPγS hydrolysis in one protomer (left). The ADP and ATPγS densities are shown in the inset.

### Mechanism of Hsp90 activation by the Aha1 co-chaperone

In the course of defining the structural mechanism by which Aha1 potently stimulates Hsp90 ATP hydrolysis, we determined 7 Hsp90:Aha1 complex structures, representing different nucleotide binding states and different domain stoichiometries. Unlike with most ATPases, in Hsp90 a large-scale conformational change rather than a chemical hydrolysis step is rate-limiting. In Hsp90 this requires breaking the Hsp90_NTD_-Hsp90_MD_ interface in the open state, closing, and reforming interactions in a rotated and Hsp90_NTD_ dimerized closed state. Notably, both Hsp90_NTD_s have to simultaneously rotate to form stabilizing interactions in the Hsp90_NTD_ dimerized closed state, making Hsp90 closure a rare event. Combining our results with previous data, we propose the following multistep model (Fig. 5) to explain how Aha1 regulates the Hsp90 conformational cycle, thereby activating hydrolysis. First, Aha1 is recruited to Hsp90 in the apo state through interactions between Aha1_NTD_ and Hsp90_MD_ as suggested by the fragment-based crystal structure^21^. Second, the binding of Aha1_CTD_ to Hsp90 induces a structure transition bringing Hsp90 from a flexible open state to a semi-closed state (Fig. 2A). In the semi-closed state, steric clashes with Aha1_CTD_ lead the Hsp90_NTD_ to undock from Hsp90_MD_ and becomes flexible. This undocking further accelerates the Hsp90 conformational cycle, because the Hsp90_NTD_ now is primed for subsequent rotation and dimerization. Third, in the presence of ATP, Aha1_CTD_ rearranges its binding interface with Hsp90_MD_ stabilizing a fully closed state which is followed by Hsp90_NTD_ dimerization (Figs. 3A-D). Fourth, the Aha1_NTD_ tilts up by 30° to interact with dimerized Hsp90_NTD_ carrying out two functions (Fig. 3F): stabilizing the Hsp90_NTD_ dimerized state and facilitating ATP hydrolysis. However, as the Aha1_NTD_ directly interacts with the lid region from only one Hsp90_NTD_, the chemical hydrolysis step is stimulated only in that protomer, leading to an asymmetric semi-hydrolyzed Hsp90 (Fig. 4B). After the hydrolysis of the second ATP, the last step is the release of Aha1 and ADP to reset Hsp90 to an open state.

**Fig. 5.**
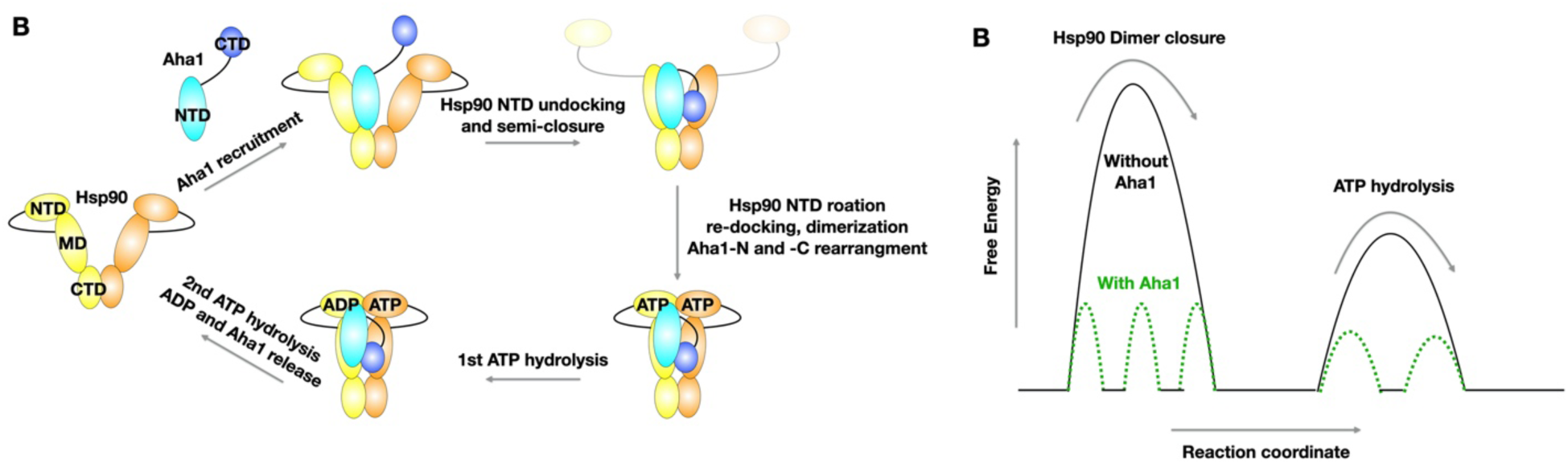
A multistep model for Hsp90 regulation by co-chaperone Aha1. (A) A 5-step model of Aha1 regulated Hsp90 conformational cycle. Hsp90 in open state recruits Aha1 through interaction between Hsp90_MD_ and Aha1_NTD_ as revealed by previous fragment-based crystal structure. The binding of full-length Aha1 induces a semi-closed Hsp90 state and undocks Hsp90_NTD_. ATP binding leads to Hsp90_NTD_ rotation, re-docking, and dimerization. The full closure of Hsp90 also accompanied by Aha1_NTD_ tilting and Aha1_CTD_ rotation. The newly established interaction between Aha1_NTD_ and Hsp90_NTD_ deterministically catalyzes the ATP hydrolysis in one of the Hsp90 protomers. The ATP hydrolysis in the second protomer results in a transient ADP-bound state followed by ADP and Aha1 release and reset the Hsp90 into the open state. (B) Aha1 functions as both a conformational and catalytic enzyme to accelerate Hsp90 dimer closure and ATP hydrolysis. It converts the large energy barrier into several smaller ones in the Hsp90 reaction pathway.

The model suggests that each Aha1 domain plays three roles in activating the Hsp90 ATPase. The Aha1_NTD_ facilitates initial Aha1 recruitment to Hsp90, stabilizes a semi-closed Hsp90 together with Aha1_CTD_, and dictates the directionality of sequential ATP hydrolysis on Hsp90. The Aha1_CTD_ on the other hand induces the semi-closure of Hsp90 in collaboration with Aha1_NTD_, undocks the Hsp90_NTD_ in the apo state, and provides an anchor point in the Hsp90 closed state so that Aha1_NTD_ can catalyze the sequential ATP hydrolysis. Much as Jencks proposed that an optimal classical enzyme takes a process with a single large energy barrier and breaks it down into multiple smaller ones, so too does Aha1 accelerate Hsp90 ATP hydrolysis. The unique feature of Aha1 is that it catalyzes transition across a large conformational barrier in addition to the more traditional high energy chemical transition state. In addition, we also discovered that it can preferentially stabilize the chemical hydrolysis step on one Hsp90 protomer, leading to an asymmetric semi-hydrolyzed intermediate state, which likely has functional consequences.

An outstanding question is what are the consequences of Aha1 acceleration of the Hsp90 conformational cycle on client protein maturation? The literature is filled with contradictory reports depending on choice of client and cell type or organism. The expression and activity of the v-Src kinase is downregulated when Aha1 is knockout in yeast^36^, consistent with Aha1 being an activator of the Hsp90 cycle. Counterintuitively, Aha1 plays an inhibitory role in CFTR folding and maturation^37^. The role of Aha1 being an inhibitor or activator can be controversial even for the same client protein, such as glucocorticoid receptor (GR). Deletion of Aha1 in yeast leads to an increase of GR expression and activity^13^. However, knockdown of Aha1 in mammalian cells does not change the GR expression but decreases hormone-dependent activation of GR^38^. These observations can perhaps be reconciled via our proposed multistep model. In addition to accelerating the overall conformational cycle, Aha1 would be expected to change the dwell time of each of the different conformational states. Thus, populations of Hsp90 states could be strongly dependent on Aha1 expression levels. Different client proteins likely utilize different Hsp90 states in distinct ways during their maturation process. For example, GR is unable to bind its ligand when bound to an open Hsp90 state but becomes active when Hsp90 closes. By contrast, a recent structure of a bound Hsp90:kinase complex revealed that the kinase becomes completely inactivated in the closed state. Making it even more complex is that Aha1 binding could occlude other cochaperone binding in a state-specific manner and this could also promote or retard client degradation. There is also the possibility of Aha1 directly competing with a possible client protein binding site, such as W296, which would be expected to hinder client loading but accelerate client release. In summary, Aha1 provides an important layer of regulation for the Hsp90 cycle and consequently Hsp90 mediated client maturation through its complicated interaction modes.

Our structures and mechanistic model also provide a framework to understand Hsp90 regulation by posttranslational modifications (PTM) and potential therapeutic intervention by small molecules. Several phosphorylation sites and SUMOylation sites have been identified on both Hsp90 and Aha1 that play important roles in regulating Hsp90 function^39–42^. A mapping of these sites onto our structures will inform on the molecular mechanism of PTM regulation. Another area of growing interest is to develop small molecules that would selectively target specific co-chaperone interactions rather than simply be ATP-competitive inhibitors on Hsp90^35,43^. This could provide the ability to manipulate only a disease-relevant subset of Hsp90-dependent cellular processes, rather than targeting all Hsp90 function which would inevitably disturb much broader cellular homeostasis.

## Acknowledgements

We thank members of the Agard Lab for helpful discussions. We gratefully thank David Bulkley, Eric Tse, Michael Braunfeld, and Glenn Gilbert from the W.M. Keck Foundation Advanced Microscopy Laboratory at the University of California, San Francisco (UCSF) for maintaining the EM facility and helping with data collection. Special thanks to Matt Harrington and Joshua Baker-Lepain for computational support on the USCF Wynton cluster. This work was supported by funding from National Institutes of Health grants U54CA209891, S10OD020054, and S10OD021741. Y.L. was supported by a Howard Hughes Medical Institute-Helen Hay Whitney Foundation Postdoctoral Fellowship, an American Heart Association Postdoctoral Fellowship, and the Program for Breakthrough Biomedical Research which is partially funded by the Sandler Foundation. M.S. was supported by 2018 AACR-Takeda Oncology Lymphoma Research Fellowship, grant number 18-40-38-SUN. D.A.A was supported by the Howard Hughes Medical Institute.

## Data availability

The electron microscopy maps and atomic models have been deposited into the Electron Microscopy Data Bank (EMDB) and the Protein Data Bank (PDB). The accession codes are EMD-22238 and PDB-6XLB for structure Hsp90^apo^-Aha1_NCNC_ (3.8 Å), EMD-22239 and PDB-6XLC for structure Hsp90^AMP^ (3.66 Å), EMD-22240 and PDB-6XLD for structure Hsp90^AMP^-Aha1_C_ (3.66 Å), EMD-22241 and PDB-6XLE for structure Hsp90^AMP^-Aha1_CC_ (2.74Å), EMD-22242 and PDB-6XLF for structure Hsp90^AMP^-Aha1_NCC_ (3.15 Å), EMD-22243 and PDB-6XLG for structure Hsp90^ATPγS^-Aha1_CC_ (2.71 Å), EMD-22244 and PDB-6XLH for structure Hsp90^ATPγS/ADP^-Aha1_NCC_ (2.83 Å), respectively.

