## Supplementary Materials for "Cryo-EM structures reveal a multistep mechanism of Hsp90 activation by co-chaperone Aha1"

This PDF file includes:

Materials and Methods
Figs. S1 to S8
Tables S1
Captions for Movies S1 to S3
References

### Materials and Methods

#### Protein cloning, expression, and purification

*S. cerevisiae* Hsc82 and Aha1 were cloned into the pET151 D-TOPO and pET28a expression plasmids, respectively, as TEV-cleavable N-terminal 6×His-tagged fusion proteins. Expression was carried out in *E. coli* BL21(DE3)-RIL. Cells were grown in TB media at 37˚C to $OD_{600}$ of ~0.8 and then induced with 0.5 mM IPTG at 16 ˚C for 18 hr. Proteins were first purified by Nickel-affinity chromatography and incubated with TEV protease overnight at 4˚C to cleave the N-terminal His-tag. After confirming the His-tag cleavage by SDS-PAGE, proteins were further purified using anion exchange and size exclusion chromatography before they were aliquoted and snap frozen with liquid $N_2$.

#### Steady-state ATPase assay

The ATPase assay monitors phosphate release via a chromogenic substrate 7-methyl-6-thioguanosine (7-MESG) in the presence of an *E. coli* PNPase (purine nucleoside phosphorylase)[1]. In these assays, 0.14 mM 7-MESG and 2 μM PNPase was used per reaction. The change in absorbance of 7-MESG was measured at 355 nm. Initial rates from phosphate release assays were obtained from fitted slopes of a linear function, $y = mx + b$, to the most linear region of each kinetic trace. The activity *versus* concentration curves were analyzed by a standard Michaelis-Menten equation.

#### Cryo-EM sample preparation

To obtain the apo state of Hsc82 in complex with Aha1, Hsc82 was incubated with Aha1 at 30 °C for 1 hour before freezing on the grid. The closed state of Hsc82 was abstained by incubating Hsc82 with 1 mM AMPPNP or ATPγS and 1 mM $MgCl_2$ at 30 °C for 30 min in the presence of various concentration of Aha1. Cryo-EM grids were prepared with Vitrobot Mark IV (FEI, Thermo Fisher Scientific, Hillsboro), using 16°C and 100% humidity. 4 μL aliquots of samples were applied to glow discharged Quantifoil R1.2/1.3, 400-mesh copper holey carbon grids (Quantifoil Micro Tools, GmbH, Großlöbichau, Germany), single blotted for 6 to 10 seconds with blot force 3, and plunge frozen in liquid ethane cooled by liquid nitrogen.

#### Cryo-EM data collection

Data were collected on a Titan Krios microscope (Thermo Fisher Scientific) operated at 300 kV with a K2 Summit direct electron detector (Gatan, Inc.). Images were recorded using SerialEM[2] with defocus varied from -0.5 to -2.7 μm for a total dose of 72 $e^{-}/Å^2$. For the datasets of Hsc82 with AMPPNP and Aha1, a super-resolution pixel size of 0.5295 Å was used and each image was dose-fractionated to 60 frames (0.2 s each, total exposure of 12 s) with a dose rate of 6 $e^{-}/Å^2/s$. For the datasets of Hsc82 with ATPγS and Aha1, a super-resolution pixel size of 0.411 Å was used and each image was dose-fractionated to 100 frames (0.1 s each, total exposure of 10 s) with a dose rate of 7.2 $e^{-}/Å^2/s$. For the datasets of apo Hsc82 with Aha1, a GIF-BioQuantum energy filter with a slit width of 20 eV was used for data collection. A super-resolution pixel size of 0.516 Å was used and each image was dose-fractionated to 100 frames (0.1 s each, total exposure of 10 s) with a dose rate of 7.2 $e^{-}/Å^2/s$. A summary of the data collection parameter was provided in Supplementary Table 1.

#### Image processing

Image stacks were motion-corrected and summed using MotionCor2[3], resulting in Fourier-cropped summed images which are binned by 2. CTFFIND4 was used to estimate defocus parameters for all the images[4]. Initial particle picking was carried out using Gautomatch without a template to generate the 2D class averages, which were then used as templates for a second-round particle picking on micrographs with 25 Å low-pass filtering. Relion 3.1[5,6] was used for all the following steps. Two rounds of reference-free 2D classification were performed for 25 iterations each with images binned by 4. Good particles were picked from 2D averages, extracted as images binned by 2, and subjected to 3D classification within Relion. An initial model was generated from crystal structure of Hsp82 (PDB: 2CG9) after deleting the co-chaperone p23 and low-pass filtered to 50 Å. Classes with the same Hsc82 and Aha1 composition were merged together, re-extracted without binning, and subjected to 3D auto-refinement. The resulting maps from refinement were post-processed and sharpened by an automatically estimated B-factor. All resolutions were estimated by applying a soft mask around the protein density and the gold-standard Fourier shell correlation (FSC) = 0.143 criterion.

**Model building, refinement and validation**

The initial model of Hsc82 was derived from the crystal structure of Hsp82 (PDB code 2CG9). There are 96.9% identity and 98.7% similarity between the two Hsp90 isoforms from *S. cerevisiae.* The initial model of yeast $Aha1_{NTD}$ were obtained from PDB code 1USV. The homology model of $Aha1_{CTD}$ was built based on NMR structure of human $Aha1_{CTD}$ (PDB code 1X53). The initial models of Hsc82:Aha1 complexes were generated by rigid-body-docking these components into the cryo-EM maps using UCSF ChimeraX[7]. The docked models were refined in real space against the cryo-EM maps using real space refinement in PHENIX[8] with secondary structure restraints, followed by iterative rounds of manual and automated refinement in Coot[9] and PHENIX[8], respectively. The final models and their fitness into the maps were visually inspected and geometry was further evaluated using MolProbity[10]. Cryo-EM data collection, refinement and validation statistics are summarized in Supplementary Table 1. Figures and Movies depicting the structures were prepared in ChimeraX[7] or VMD[11].

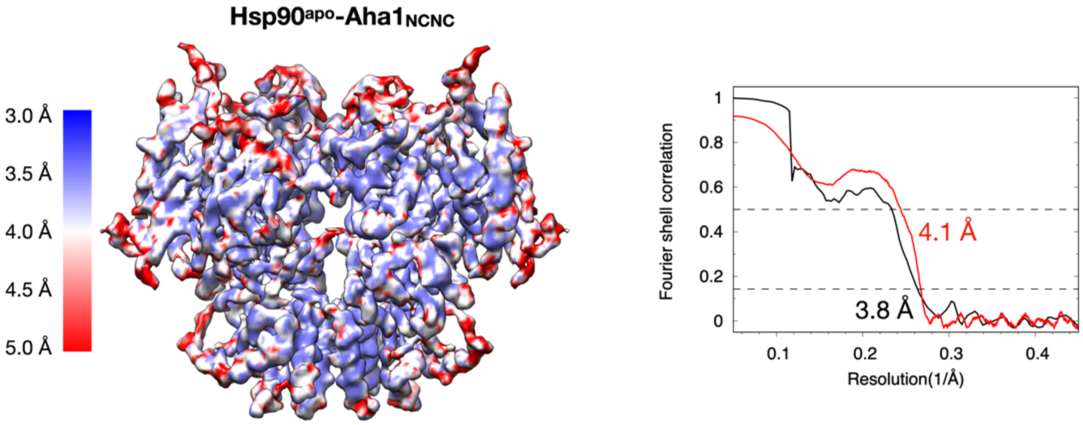


**Fig. S1.** Local resolution map and gold-standard Fourier shell correlation (FSC) curves for Cryo-EM structure of Hsp90$^{apo}$-Aha1$_{NCNC}$. Note the Hsp90$_{NTD}$s are completely missing.

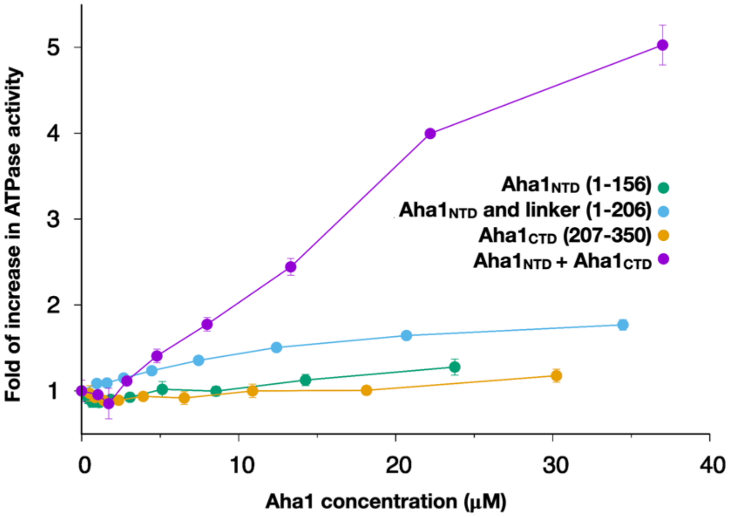


**Fig. S2.** Aha1 truncations significantly reduce its accelerating effect on the rate of Hsp90 ATP hydrolysis. In the context of the Aha1$_{NTD}$ the linker provides additional stimulation. The two domains can work together even in the absence of covalent linkage (Aha1$_{NTD}$+Aha1$_{CTD}$), but the level of stimulation is 6-fold lower than the full-length Aha1.

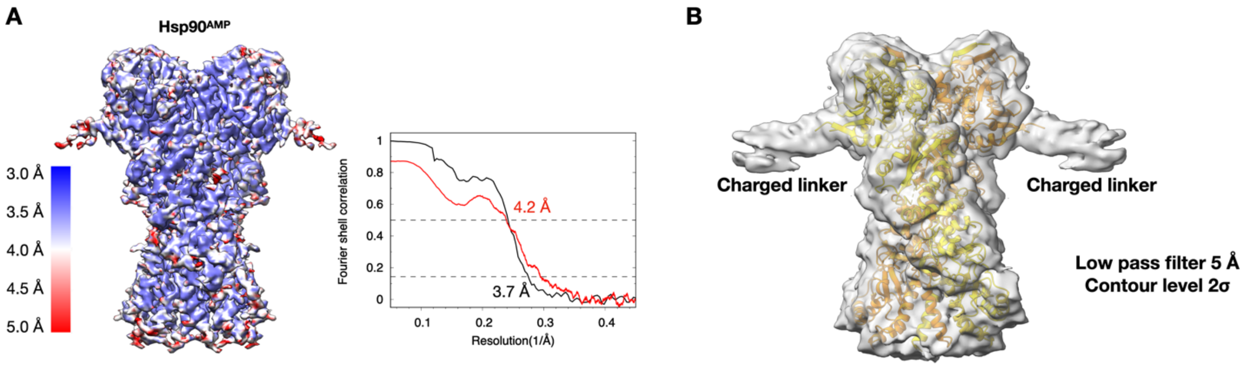


**Fig. S3.** Cryo-EM structure of Hsp90$^{AMP}$. (A) Local resolution map and gold-standard Fourier shell correlation (FSC) curves. (B) Visualization of the charged linker region by low pass filtering the map to 5 Å.

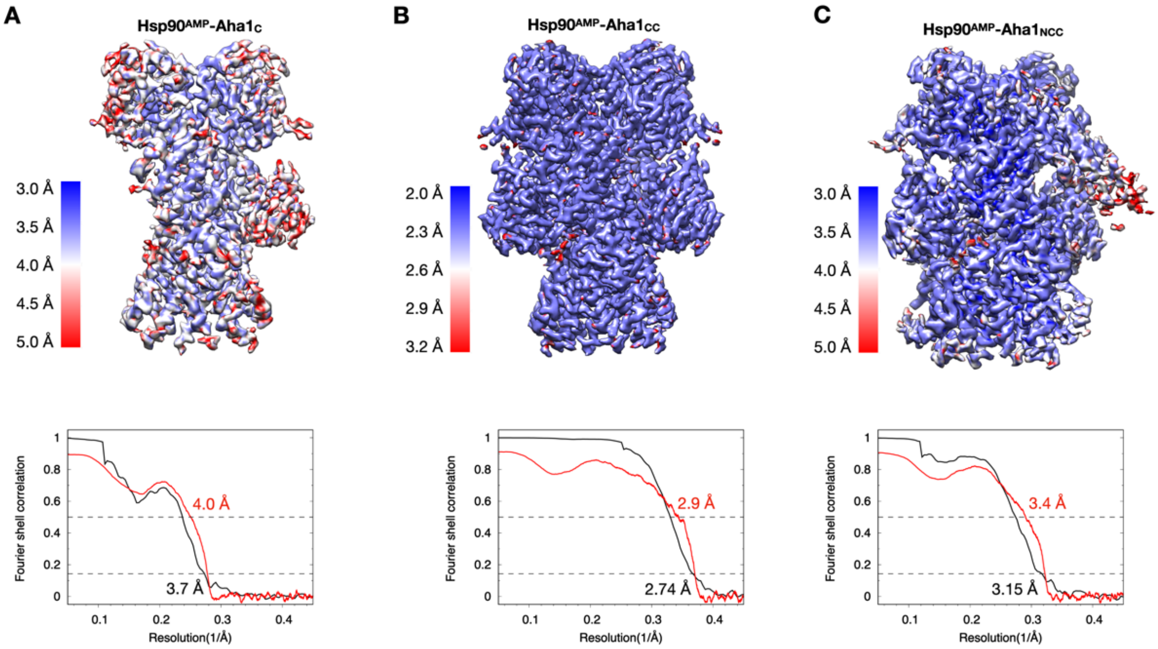


**Fig. S4.** Cryo-EM structures of Hsp90:Aha1 complexes in the presence of AMPPNP. (A) Local resolution map and gold-standard Fourier shell correlation (FSC) curves for cryo-EM structure of Hsp90$^{AMP}$-Aha1$_{C}$. (B) Local resolution map and gold-standard Fourier shell correlation (FSC) curves for cryo-EM structure of Hsp90$^{AMP}$-Aha1$_{CC}$. (C) Local resolution map and gold-standard Fourier shell correlation (FSC) curves for cryo-EM structure of Hsp90$^{AMP}$-Aha1$_{NCC}$.

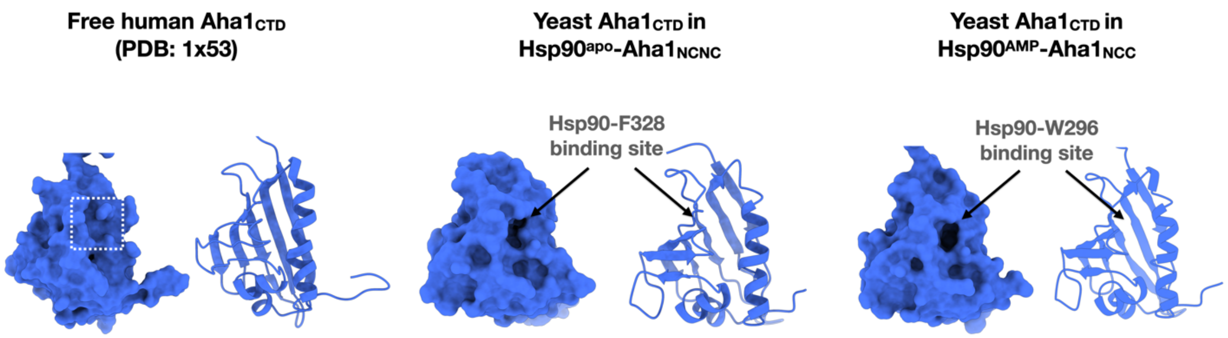


**Fig. S5.** Hydrophobic pocket formed on the surface of Aha1 when bound to Hsp90. The size and depth of the pocket increase in response to either F328 in $Hsp90^{apo}$ and even further as induced by W296 in the closed Hsp90.

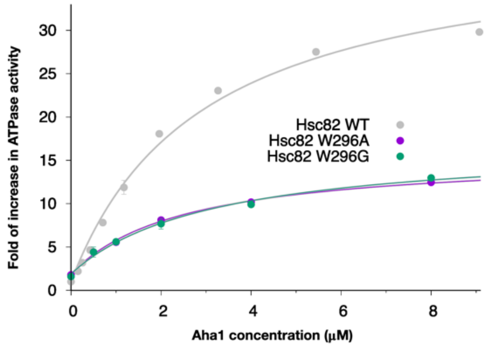


**Fig. S6.** Hsp90 W296A and W296G mutations reduce the Aha1 activating effect by 6-fold.

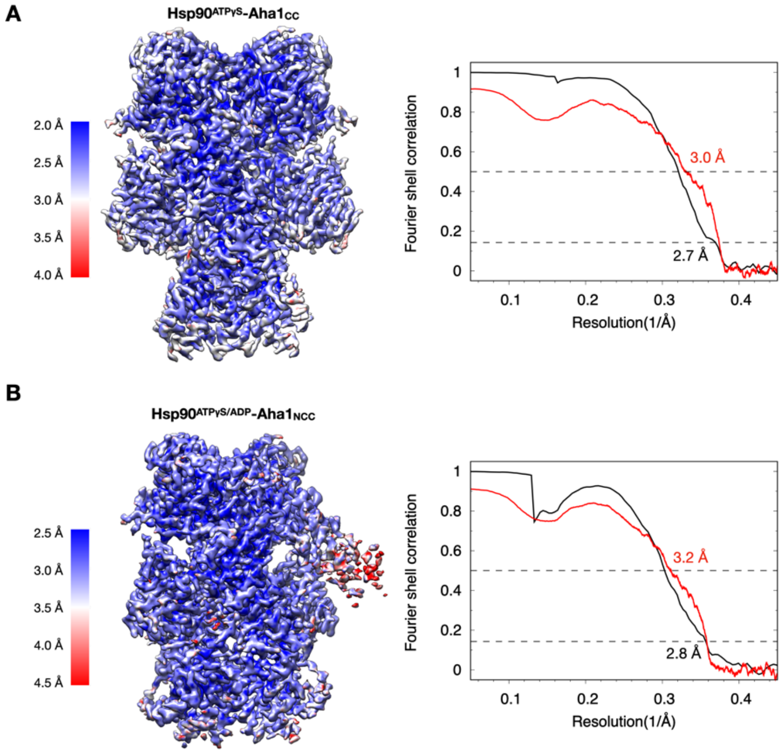


**Fig. S7.** Cryo-EM structures of Hsp90:Aha1 complexes in the presence of ATPγS. (A) Local resolution map and gold-standard Fourier shell correlation (FSC) curves for cryo-EM structure of Hsp90 $^{ATP\gamma S}$-Aha1$_{CC}$. (B) Local resolution map and gold-standard Fourier shell correlation (FSC) curves for cryo-EM structure of Hsp90$^{ATP\gamma S/ADP}$-Aha1$_{NCC}$.

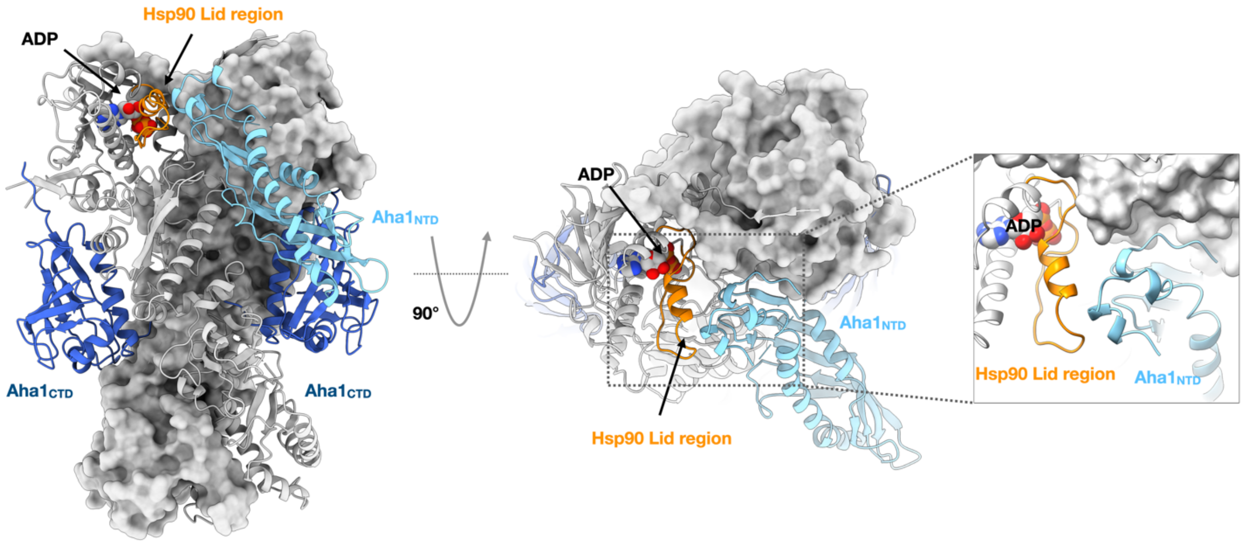


**Fig. S8.** Aha1$_{NTD}$ forms close contact with the Hsp90 lid region in the presence of nucleotides and facilitates the ATP hydrolysis.

**Table S1.** Cryo-EM data collection, refinement and validation statistics

| | #1 $Hsc82^{apo}$-$Aha1_{NCNC}$ (EMD-22238) (PDB 6XLB) | #2 $Hsc82^{AMP}$ (EMD-22239) (PDB 6XLC) | #3 $Hsc82^{AMP}$-$Aha1_{C}$ (EMD-22240) (PDB 6XLD) | #4 $Hsc82^{AMP}$-$Aha1_{CC}$ (EMD-22241) (PDB 6XLE) | #5 $Hsc82^{AMP}$-$Aha1_{NCC}$ (EMD-22242) (PDB 6XLF) | #6 $Hsc82^{ATP\gamma S}$-$Aha1_{CC}$ (EMD-22243) (PDB 6XLG) | #7 $Hsc82^{ATP\gamma S/ADP}$-$Aha1_{NCC}$ (EMD-22244) (PDB 6XLH) |
|---|---|---|---|---|---|---|---|
| **Data collection and processing** | | | | | | | |
| Microscope | FEI Titan Krios | FEI Titan Krios | FEI Titan Krios | FEI Titan Krios | FEI Titan Krios | FEI Titan Krios | FEI Titan Krios |
| Camera | Gatan K2 Summit | Gatan K2 Summit | Gatan K2 Summit | Gatan K2 Summit | Gatan K2 Summit | Gatan K2 Summit | Gatan K2 Summit |
| Energy Filter | 20 eV slit | n/a | n/a | n/a | n/a | n/a | n/a |
| Magnification | 130,000x | 22,500x | 22,500x | 22,500x | 22,500x | 29,000x | 29,000x |
| Voltage (kV) | 300 | 300 | 300 | 300 | 300 | 300 | 300 |
| Electron exposure (e–/Å$^2$) | 72 | 72 | 72 | 72 | 72 | 72 | 72 |
| Defocus range (μm) | -0.6 to -2.7 | -0.8 to -2.0 | -0.6 to -2.0 | -0.5 to -2.0 | -0.5 to -2.0 | -0.6 to -2.0 | -0.6 to -2.0 |
| Pixel size (Å) | 1.032 | 1.059 | 1.059 | 1.059 | 1.059 | 0.822 | 0.822 |
| Symmetry imposed | C2 | C2 | C1 | C2 | C1 | C2 | C1 |
| Micrographs (no.) | 2,765 | 2,555 | 10,364 | 10,364 | 4741 | 4293 | 4293 |
| Initial particle images (no.) | 835,494 | 719,342 | 3,184,838 | 3,184,838 | 2,544,160 | 1,143,371 | 1,143,371 |
| Final particle images (no.) | 62,906 | 48,464 | 32,301 | 399,013 | 76,347 | 126,867 | 112,624 |
| Map resolution (Å) FSC threshold 0.143 | 3.8 | 3.66 | 3.66 | 2.74 | 3.15 | 2.71 | 2.83 |
| Map resolution range (Å) | 3.3-6.3 | 3.3-6.3 | 3.3-5.8 | 2.3-3.8 | 2.8-5.8 | 1.8-3.8 | 2.3-5.1 |
| **Refinement** | | | | | | | |
| Initial model used (PDB code) | 2CG9,1USV,1X53 | 2CG9 | 2CG9, 1USV | 2CG9, 1USV | 2CG9,1USV,1X53 | 2CG9, 1USV | 2CG9,1USV,1X53 |
| Model resolution (Å) FSC threshold 0.5 | 4.1 | 4.2 | 3.9 | 3.0 | 3.4 | 3.0 | 3.2 |
| Map sharpening *B* factor (Å$^2$) | -107 | -115 | -111 | -107 | -87 | -90 | -75 |
| Model composition | | | | | | | |
| Non-hydrogen atoms | 11184 | 9864 | 10997 | 12266 | 13660 | 12266 | 13617 |
| Protein residues | 1380 | 1212 | 1349 | 1502 | 1680 | 1504 | 1675 |
| Ligands | 0 | 2 | 2 | 2 | 2 | 2 | 2 |
| Ions | 0 | 2 | 2 | 4 | 4 | 4 | 4 |
| *B* factors (Å$^2$) | | | | | | | |
| Protein | 93.22 | 43.76 | 61.28 | 43.19 | 57.68 | 35.31 | 47.02 |
| Ligand | n/a | 30.06 | 47.84 | 23.35 | 39.68 | 17.77 | 40.79 |
| R.m.s. deviations | | | | | | | |
| Bond lengths (Å) | 0.001 | 0.001 | 0.001 | 0.001 | 0.001 | 0.001 | 0.001 |
| Bond angles (°) | 0.416 | 0.359 | 0.355 | 0.347 | 0.342 | 0.339 | 0.373 |
| Validation | | | | | | | |
| MolProbity score | 1.40 | 1.33 | 1.31 | 1.27 | 1.40 | 1.32 | 1.33 |
| Clashscore | 7.28 | 6.07 | 5.72 | 5.07 | 6.23 | 5.84 | 5.95 |
| Poor rotamers (%) | 0.4 | 0.00 | 0.70 | 0.60 | 0.60 | 0.00 | 0.13 |
| Ramachandran plot | | | | | | | |
| Favored (%) | 98.03 | 98.41 | 98.5 | 98.58 | 98.37 | 98.65 | 98.49 |
| Allowed (%) | 1.97 | 1.59 | 1.5 | 1.42 | 1.63 | 1.35 | 1.51 |
| Disallowed (%) | 0.00 | 0.00 | 0.00 | 0.00 | 0.00 | 0.00 | 0.00 |

**Movie S1.** Hsp90 structural transition from open to semi-closed state in the absence of nucleotide, which is induced by Aha1 binding. In the presence of nucleotides, Hsp90 transitions from a semi-closed state to fully closed state. Aha1 and $Hsp90_{NTD}$ were not shown from clarity purpose.

**Movie S2.** $Aha1_{NTD}$ tilt up 30º when Hsp90 transitions from the apo semi-closed state to nucleotide bound fully closed state.

**Movie S3.** Aha1 structural transition when Hsp90 changes from the apo semi-closed state to nucleotide bound fully closed state. $Hsp90_{NTD}$s were not shown for clarity purposes.
